# Network Ontology Transcript Annotation Identifies Genetic Signals Underlying Sex Determination

**DOI:** 10.1101/2025.05.05.650505

**Authors:** Leonardo R. Orozco, Audrey E. Weaver, Daniel J. Klee, Christopher S. Pauli, Christopher J. Grassa, Daniela Vergara, Anthony Baptista, Kristin White, Benjamin F. Emery, Natalie R.M. Castro, Shiva Garuda, Rafael F. Guerrero, Brian C. Keegan, Nolan C. Kane

## Abstract

*Cannabis sativa* L. (marijuana, hemp, cannabis) is an angiosperm species currently evolving sex chromosomes. Genetic mechanisms, primarily an XY chromosome system, dictate cannabis sex expression in dioecious populations. However, sexual expression is also governed by the interplay of hormone regulatory gene networks, influenced by both genetic and environmental factors. Within the species, some populations exhibit dioecy, monoecy, or a gradient of both. Dioecious individuals produce exclusively male or female flowers, while monoecious plants bear both male and female flowers. Remarkably, through interruption of phytohormone signal transduction via abiotic stressors, genetically male or female cannabis are able to produce flowers of the opposite sex. Previous transcriptomic analysis have identified genes associated with masculinization through the application of phytohormone signal disruption using silver thiosulfate treatment. We analyzed transcriptomic data from cannabis treated with colloidal silver to similarly induce masculinization. Using Nota (Network Ontology Transcript Annotation), a multilayer network analysis (random walk with restart) tool, we identified candidate genes involved in sex-determination. Nota and a companion program Jack facilitate multi-layer network analyses, enabling discovery and annotation of gene-trait associations. Our findings highlight Nota’s robust application to enrich the genetic architecture of complex traits, particularly in non-model systems like cannabis, and complex traits such as sex determination. Our analyses identified genes associated with cell wall morphogenesis and embryogenic tissue homeostasis, indicating that silver ion treatment perturbs phytohormone signal transduction through metal ion imbalance. In this reproductive strategy, cannabis is able to make use of its widely investigated sex-determining genetic architecture to navigate transient changes in co-expression and cross-cellular signaling driving embryogenic cell wall re-patterning.

## Introduction

*Cannabis sativa* L. (cannabis, hemp, marijuana, hereafter “cannabis”) has long been a controversial species with an equally complex domestication^1^. Due to it’s historically polarizing social context, and divergent human breeding objectives—food, fiber, medicinal and psychoactive traits^2–4^—there is a high degree of genetic diversity observed in the species. This genetic landscape reflects the irregularity of its domestication, shaped by both illicit and regulated breeding efforts. Government-produced and regulated accessions show significantly reduced genetic diversity compared to those in unregulated markets, exemplifying how inconsistent breeding practices have shaped the current gene pool^5^. Cannabis has a diploid genome (2n=20) characterized by nine autosomes and a pair of sex chromosomes, with males, the heterogametic sex, harboring X and Y chromosomes and females carrying two X chromosomes^6^. Furthermore, genes related to sex expression are dispersed throughout the genome, with roughly 74% found on autosomes, indicating that the regulation of sexual traits is widespread and not restricted solely to the sex chromosomes^7–9^.

Different populations of cannabis have distinct reproductive mating strategies^1,10^: plants can be dioecious, with individuals of distinct male and female sexes, or monoecious, with both reproductive organs occurring on the same individual. Dioecious female individuals(XX) are the main source of cannabinoids for medicinal and recreational purposes. Cannabinoid and terpenoids are produced and stored in glandular trichomes, the secretory organs concentrated in female inflorescence tissue^11^. Monoecious individuals are prioritized for fiber and grain varieties, and low cannabinoid and terpene biosynthesis^12^. Monoecious individuals also appear to have two X chromosomes^13,14^, yet exhibit sex expression plasticity that transcends chromosomal expectation^15^, highlighting the dynamic regulation of sexual phenotype beyond karyotype alone. Un-pollinated females, deprived of the energetic cost of seed production, maintain an enhanced capacity for secondary metabolism^16,17^. The induction of increased trichome production in un-pollinated female flowers was likely originally an adaptive strategy to increase defense of reproductive tissue as seen in other secondary trichome holding plants^18^. Trichome abundance is a priority for in breeding programs prioritizing high secondary metabolite yielding flower tissue. Selective breeding goals in cannabis are challenged by environmental stress and their disruption of steady state metabolism which governs the life history strategy of individual plants^10^.

Plants are sophisticated biological systems that have evolved to maintain physiological flexibility while undergoing environmental change^19^. This physiological flexibility allows plants to adjust developmental and metabolic processes to preserve reproductive success across a range of stress conditions^20^. In this context, stress can be broadly defined as any environmental signal that alters plant physiology^21^. Glandular trichomes mediate metabolic and stress signaling disruptions of drought, ultraviolet radiation, pests and the absence of pollen during flowering periods^17^. They have also been reported to assist in extracellular storage and transportation of critical metabolites across membranes^18,22^. The metabolism of glandular trichomes is highly responsive to environmental signaling cues^23^. This responsiveness possibly enables cannabis to differentiate reproductive tissue development strategy in response to environmental stress signals. Similar relationships between glandular trichomes and epidermal development in tomato fruiting tissue suggests a shared developmental framework among plant reproductive tissues, wherein trichomes integrate environmental cues into their growth and differentiation^18,24^.

Environmental stress signals directly contribute to sex determination in cannabis^7,15,25^. Factors such as temperature variations, photoperiod, and the administration of hormones or chemicals interacting with plant hormonal pathways can all influence the reproductive development of cannabis^26^. The interplay between genetic expression and environmental constraints represent a dynamic system where the plant adjusts gene co-expression to secure reproductive output. Plant hormones (phytohormones) are transient regulators in this cross-cellular communication^27–29^. Ethylene in particular, orchestrates processes including fruit ripening, shoot development, germination, nodulation, and floral senescence^30^. Exogenous application of growth promoting ethephon, a synthetic ethylene-releasing compound, can rewire phytohormone signaling to prioritize the release of ethylene^31^. This uptick in ethylene availability can induce feminization in cannabis^32^.

Conversely, silver-based treatments disrupt internal ethylene signaling by binding or out-competing endogenous metal ions required for its synthesis. This heavy metal ion disruption of ethylene transcription and biosynthesis has been observed to induce a male-forward reproductive strategy in cannabis^7,33–35^. These treatments do not rewrite or alter genomic architecture; they alter the expression patterns of sex-determining genes^7,9,25,36^. These metal-induced male flowers are able to pollinate and thus yield “feminized seed” lacking a y chromosome^37–39^. These chemical interventions thus serve as experimental changes, revealing the genetic plasticity cannabis can employ to reorganize it’s reproductive development in response to heavy-metal interruption of ethylene transcription using alternative phytohormone signal transduction.

To characterize alternative signaling transduction routes in the cannabis sex-determining cellular pathways, we followed and extended the methods of Zhu et al. 2023, who predicted drought stress-related genes in rice using a RWR analysis on a multilayer biological network^40^. Recently, multilayer networks have become powerful frameworks for modeling organisms comprised of multiple biological layers^41^. A multilayer biological network represents different types of relationships among and across the network members (genes and proteins), this in part is made possible via network-connecting bipartite networks. Connecting different biological layers from the same experiment enables integrated modeling of transcriptional coordination and macromolecular interaction. Random walk (RW) is a novel approach to exploring the structure and connection of biological networks.

RW was inspired by the PageRank algorithm^42^, which was initially developed for ranking web pages in search results by simulating the behavior of an internet user following hyperlinks or restarting on arbitrary pages. In a biological context by simulating the movement of a network walker randomly traversing biological connections, random walk algorithms are able to capture several structural properties of biological networks^43^, including connectivity^44^, community structure^45^, and centrality^46^ of network members. Random Walk with Restart (RWR) an extension of RW was first introduced by Pan et al. 2004^47^. In RWR the network walker, at each step, can navigate from one network member to one of its neighbors or restart its walk from a node randomly sampled from a set of network members referred to as seed nodes. By enabling restart from one or several seed nodes, RWR simulates a diffusion process in which the objective is to determine the steady state of an initial probability distribution^48^. This steady state represents a measure of proximity between the seed(s) and all the network members, rather than a centrality measure as in PageRank.

Here we present **Nota** (Network Ontology Transcript Annotation), a multilayer network analysis bioinformatic tool. Nota conducts a Random Walk with Restart (RWR) algorithm adapted from `multixrank` a Python package^49,50^. Using differentially expressed genes as seed nodes (where Nota starts and restart its’ walk) to infer additional related genes based on their proximity and the strength of their connection. By diffusing probability of network member relatedness across the multilayer networks, Nota’s RWR quantifies the extent to which influence from differentially expressed genes spreads through the multilayer biological network. This cross network probability diffusion reveals which genes and their encoded proteins were most responsive to experimental conditions in transcriptomic studies.

To investigate the cross-cellular signaling mechanisms enabling the plasticity of sex determination in cannabis, we employed Nota and Jack to curate and analyze a multilayer biological network composed of a gene co-expression layer, a species-representative proteinprotein interaction (PPI) layer^51^, and a bipartite gene-to-protein annotation layer linking them in a 1:1 relationship. We performed differential expression analysis to identify statistically significant transcriptional differences among male, female, and colloidal silverinduced male flowers grown under identical environmental conditions(Figure 1). Genes associated with phytohormone signaling, metal-ion homeostasis, lipid metabolism, cell-wall remodeling, transcriptional regulation, stress response, and both primary and secondary metabolism were examined, and Nota was used to infer additional gene associations from these contrasts. Our analysis is guided by the hypothesis that the high metabolic cost of cannabis masculinization are rebalanced through alternative transcriptional and phytohormone signal-transduction mechanisms that restore developmental physiology to a steady metabolic state under environmental stress. We further hypothesize that cross-talk among phytohormone signaling networks regulating stress response, development, and embryogenic cell-wall re-patterning underlies the plasticity of sexual determination in cannabis.

**Figure 1.**
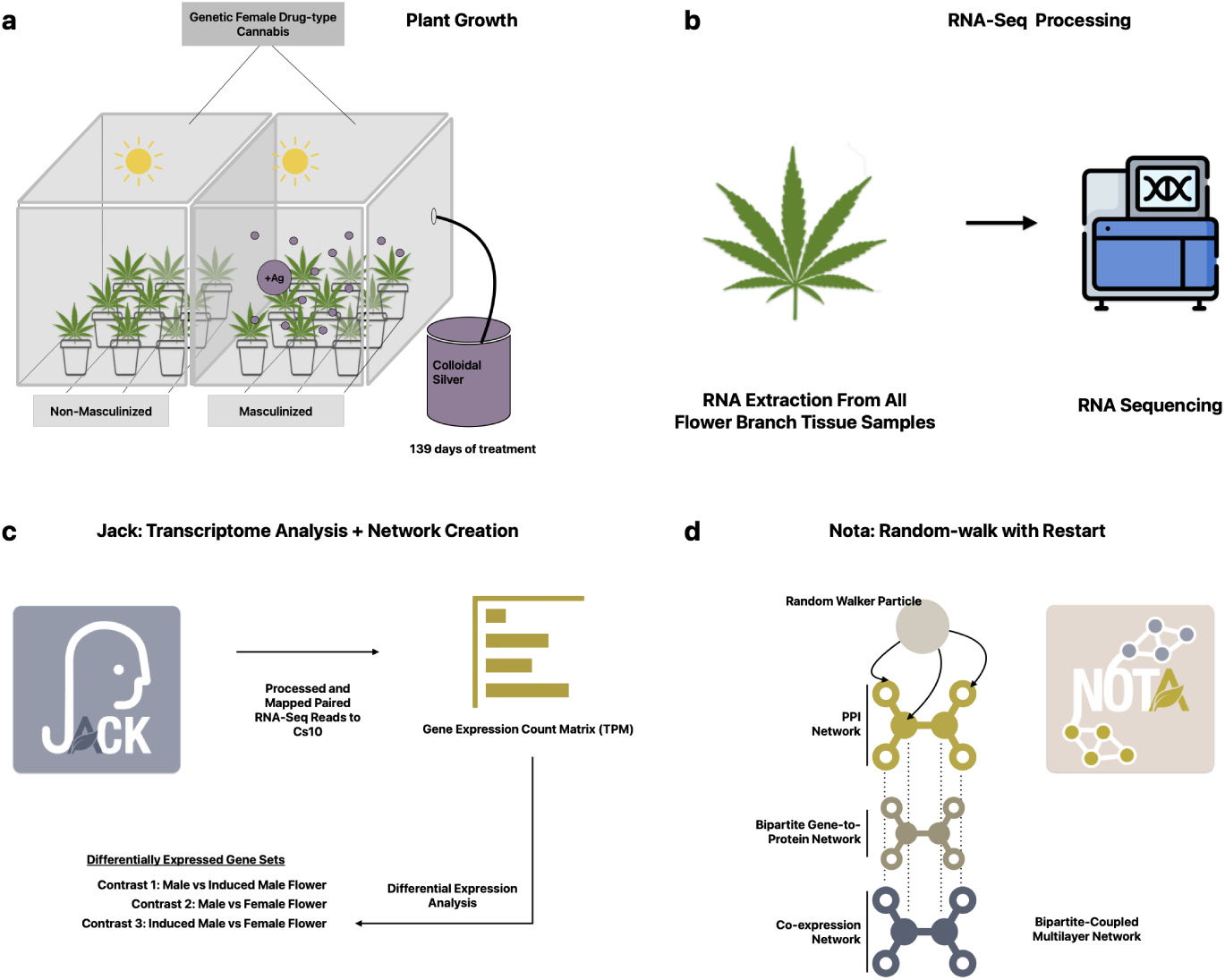
Experimental Work Flow. (a) Plant Growth and Treatment: Genetically Female cannabis plants were grown under standard conditions. A subset of plants was treated with colloidal silver to induce masculinization in selected branches. (b) RNA Sequencing: RNA was extracted and sequenced from male flower (MF), female flower (FF), and induced male flower (IMF) after 139 days of treatment. (c) Bioinformatic and Statistical Analysis: Raw RNA-seq reads were processed and mapped to the cannabis reference genome (Cs10) using Jack. Differential expression analysis was performed using DESeq2 to identify genes differentially expressed between the three groups (MFvsIMF, MFvsFF, and IMFvsFF). (d) Nota: Network Ontology Transcript Annotation, a tool employing a specialized random walk with restart algorithm, was applied to the network to identify key proteins and pathways associated with sexual plasticity.

## Results

### Differential Expression Analysis

We constructed a DESeq2 object, to compare our true male flower (MF), true female flower (FF), and induced male flower (IMF) RNA-seq samples (Orozco dataset). We parsed and filtered (see Differential Expression Analysis subsection in Methods) differentially expressed genes (DEGs) from the tabulated result object for each treatment contrast, MFvsIMF, MFvsFF, and IMFvsFF. Thresholding for DEGs was set to an absolute log_2_fold change value greater than or equal to 1 (log_2_FC |1|) and an adjusted p value of less than or equal to 0.05 (p-adj 0.05). The MFvsFF contrast was used as a control for comparison of the MFvsIMF and IMFvsFF differential expression data. This analysis was replicated on the RNA-seq samples from Adal et al. (2021), in parallel using identical methods and parameters.

For the Orozco dataset, the differential expression analysis yielded 3,324 DEGs in MFvsIMF, 3,504 DEGs in MFvsFF, and 133 DEGs in IMFvsFF (Supplementary Table S1 online). In these respective sets of DEGs, there were a number of genes significantly differentially expressed in two or more contrasts. 81.3% (2,728) of DEGs in MFvsIMF were also differentially expressed exclusively in MFvsFF. In comparison, 63.9% (85) of DEGs in the IMFvsFF contrast were also exclusively found in MFvsFF. Additionally, only 0.2% of DEGs in MFvsIMF were also exclusively differentially expressed in IMFvsFF. Lastly, there were 26 genes identified as differentially expressed in all three contrasts (Figure 2). Full overlapping DEG lists and their encoded proteins can be seen in Orozco Contrast Overlap DEGs with Proteins supplemental data folder online.

**Figure 2.**
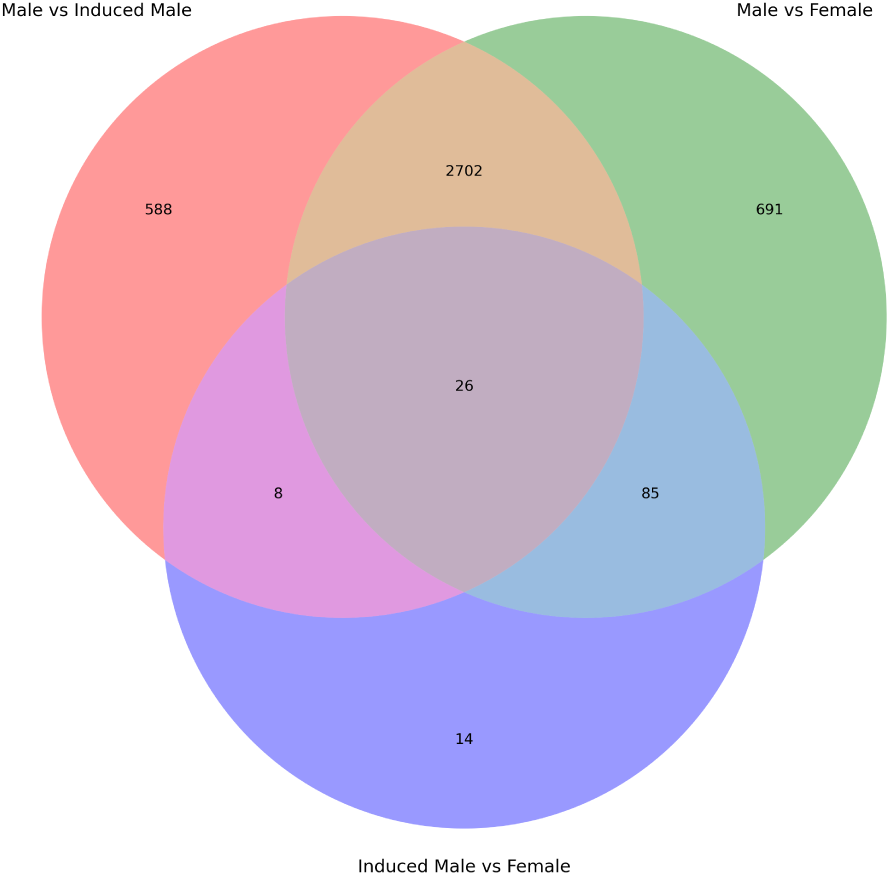
Orozco dataset overlapping DEGs by contrast Venn Diagram displaying the number of DEGs (log_2_FC |1| and p-adj 0.05) found in each respective contrast as well as overlapping between two or all contrasts within the Orozco RNA-seq samples. Full overlapping DEG lists and their encoded proteins can be seen in Orozco Contrast Overlap DEGs with Proteins supplemental data folder online.

From the analysis of the Adal et al. (2021) dataset, we identified: 2,247 DEGs in MFvsIMF, 20,014 DEGs in MFvsFF, and 2,433 DEGs in IMFvsFF (Supplementary Table S2 online). Further identification of overlapping sets of up-regulated (log_2_FC 1) and down-regulated (log_2_FC -1) DEGs across comparable pairwise contrasts in each respective dataset was done (Supplementary Table S3 online). Direction of regulation corresponds to the first referred to treatment’s alteration of transcription i.e when discussing down-regulation in the MFvsIMF contrast, we will be referring to genes that are down-regulated in the MF treatment compared to the IMF treatment. In the MFvsIMF contrast, 76.2% of DEGs (2,534) were down-regulated in the Orozco samples while only 22% of DEGs (494) were down-regulated in the Adal samples. In the MFvsIMF contrast, we identified 69 up-regulated genes and 56 down-regulated genes overlapping between both sample sets. For the MFvsFF contrast, 29.2% (1,022) of DEGs in the Orozco samples were up-regulated whereas 99.8% (19,973) of DEGs in the Adal samples were up-regulated. Of the respective MFvsFF DEGs, there were 1,002 consistently up-regulated and 6 down-regulated DEGs overlapping between the two sample sets. Lastly, for the IMFvsFF contrast, 92.5% (123) of DEGs were up-regulated in the Orozco samples and 30.6% (744) of DEGs were up-regulated in the Adal samples. In the IMFvsFF contrast, there was only 1 consistently up-regulated gene (LOC11570581: pathogenesis-related protein) across both datasets and 0 consistently down-regulated genes. See full DEG lists for each contrast in Merged Datasets DEGs supplementary data folder online.

### Random-Walk-with-Restart Inference

To infer additional sex determination-related genes, we next performed a Random-Walk-with-Restart (RWR) analysis. Using Nota, we conducted a personalized RWR analysis on our bipartite-coupled multilayer network, employing three sets of seed nodes derived from DEGs found in each respective experimental contrast. Our network inference identified additional candidate genes involved in the genetic architecture underlying masculinization in female cannabis treated with colloidal silver. The RWR algorithm calculates the geometric mean for each seed set, ensuring that the resulting scores and inferences sum to 1. Here, we report the topologically related gene inferences for each contrast, excluding seed nodes and genes with a geometric mean score below our confidence thresholds (MFvsIMF: 157, MFvsFF: 184, IMFvsFF: 236). Of these inferences, some overlapped when checked against our DEG lists (MFvsIMF: 42, MFvsFF: 77, IMFvsFF: 23) (See full inference lists in RWR Results and Multiplex Files supplemental data folder online). The presence of overlap of inferred genes and DEGs validated the RWR algorithm’s ability to infer candidate genes as those were not provided as seed nodes. All inferred genes were expressed in the combined Orozco and Adal sample sets used to construct the co-expression network. We generated four ontology-enriched PPI networks, including one comprising inferred sex determination-related genes and their encoded proteins shared across all three contrast inference sets. The other three PPI networks were separate ontology reports for each contrast’s inferences and were made by selecting them as local cluster networks within our species representative cannabis PPI network available on StringDB^51^.

### Gene Network Ontology of Shared RWR Inferences

We inferred a total of 577 genes with confident geometric mean scores, of which 433 were shared across all three inference sets (Figure 3). Our shared network cluster consists of 433 nodes and 470 edges, with an average node degree of 2.17 and a local clustering coefficient of 0.287, with high-confidence interaction scores (>0.9). The overall PPI enrichment was highly significant (p-value < 1.0e-16). We then applied k-means clustering to group functionally related proteins, ensuring that all disconnected sub-graphs were accounted for. This PPI network shows the most biologically relevant ontology profiles of the inferences shared across all contrasts.

**Figure 3.**
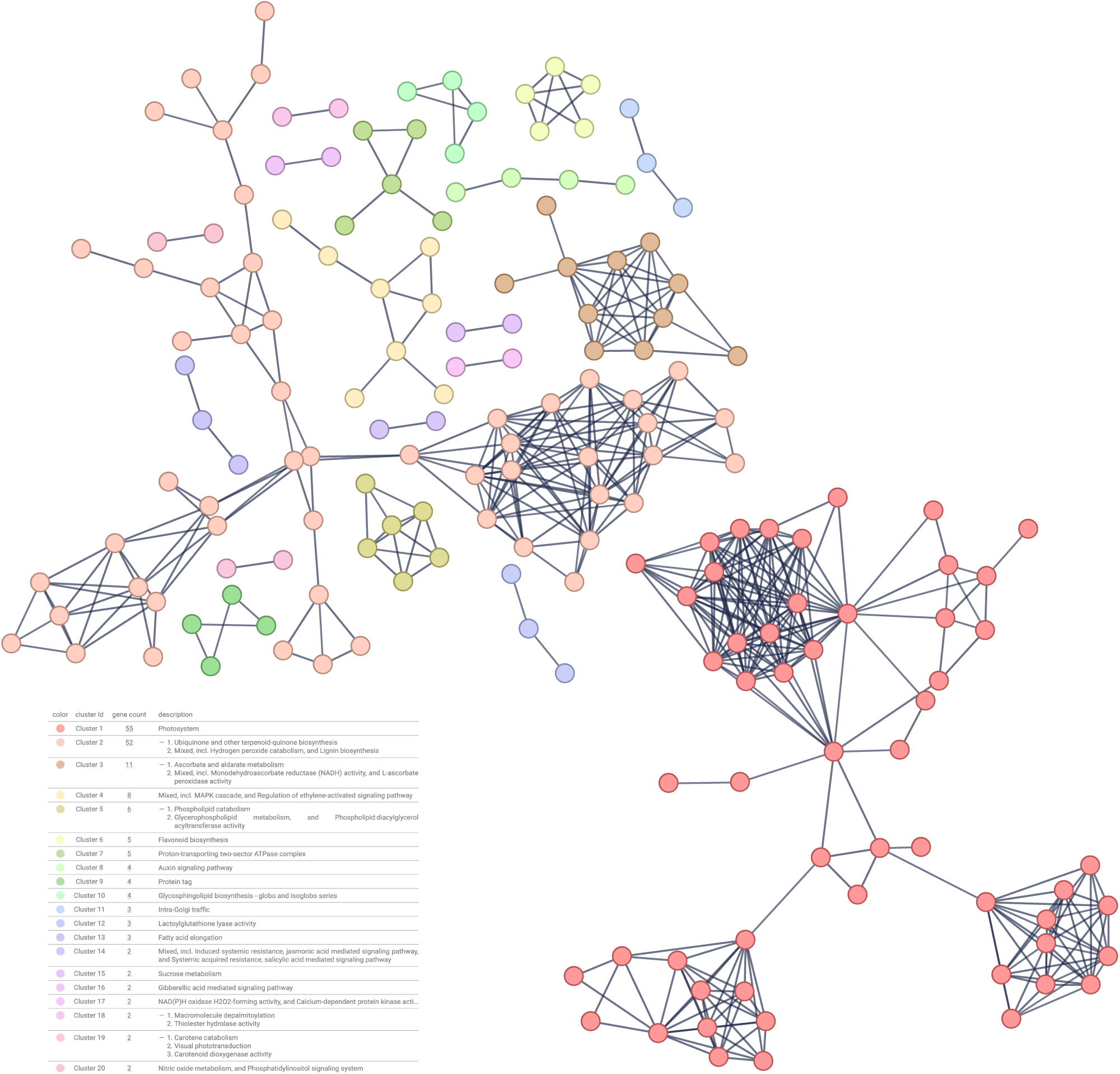
K-Mean 20 clustering of Shared Non-unique Inferences. This network was made by highlighting all 577 inferred genes to to our species representative PPI. We selected proteins with high-confidence (>/=0.9) interactions shared between them, post this filtering we clustered the 433 non-unique genes. We hid all the non-connected nodes and removed node labels for visual clarity. Edge color opacity is correlated with interaction scores (darker correlates to higher scoring interactions).

This unique set of inferences exhibited gene ontology (GO) enrichment for biological processes related to photosynthesis, spliceosomal snRNP assembly, cellular component assembly involved in morphogenesis, and secondary metabolism. Additionally, we identified enrichment for molecular functions associated with ligase and oxidoreductase activity. Consistently, many enriched cellular component terms were linked to photosystem I and photosystem II, the mediator complex, the SMN-Sm protein, and spliceosome complexes. Several genes were associated with both primary metabolism (e.g., pyruvate, tyrosine, cysteine, and methionine metabolism) and secondary metabolic pathways, including ubiquinone and other terpenoid-quinone biosynthesis, ascorbate and aldarate metabolism, pyruvate metabolism, and tyrosine metabolism (KEGG,^52^). Furthermore, we observed enrichment in reactome pathways such as fatty acid metabolism and signaling by receptor tyrosine kinases.

### Male Flower vs Induced Male Flower Inferred Gene Network Topology

The network consisted of 157 genes, 224 edges, an average node degree of 2.85, an average local clustering coefficient of 0.324, and an overall PPI enrichment p-value of < 1.0e-16. These genes were inferred to be related to the genes of interest found in differential expression results from the MFvsIMF contrast. Following the filter of proteins sharing high confidence interaction scores, we ran k-means clustering (accounting for the minimum disconnected graphs in the network) and identified 10 clusters (see corresponding subsection supplemental note for full cluster associations). Cluster 1 was the largest (22 genes) with enrichment in secondary metabolic pathways such as phenylpropanoid metabolism and the remodeling of diacylglycerol (DAG) and triacylglycerol (TAG) as well as ubiquinone and other-terpenoid biosynthesis. We also observed clusters related to photosystems components, redox homeostasis (ascorbate and aldarate metabolism, NADH-dependent reductase activity), and sucrose metabolism. Additionally, smaller clusters were enriched in hormonal and stress response pathways (Induced systemic resistance, jasmonic acid and salicylic acid signaling pathways, Regulation of jasmonic acid signaling and ubiquitin-like conjugation), while two unannotated proteins (XP_030490813.2 and XP_030493450.2) were found to be related to lipid metabolism.

### Male Flower vs Female Flower Inferred Gene Network Topology

This high-confidence interaction network consisted of; 184 genes, 198 edges, an average node degree of 2.15, an average local clustering coefficient of 0.334, and an overall PPI enrichment p-value of < 1.0e-16. Following the same procedure for the MFvsIMF contrast inferences we identified 14 functional clusters (see corresponding subsection supplemental note for full cluster associations). The largest clusters were enriched for transcriptional regulation and lipid-associated metabolism, including genes encoding mediator complex subunits, enzymes involved in phenylpropanoid and vitamin E biosynthesis, ubiquinone and terpenoidquinone pathways, and multiple acyl-transferase and phospholipid-modifying activities. Additional clusters were associated with energy and redox metabolism, encompassing mitochondrial ATP synthase components, fatty acid elongation, and flavonoid biosynthesis. Smaller clusters contained genes linked to protein tagging, glycosphingolipid biosynthesis, and intra-Golgi trafficking, along with two unannotated proteins associated with glucose metabolic processes.

### Induced Male Flower vs Female Flower Inferred Gene Network Topology

This inferred gene network consisted of; 236 genes, 85 edges, an average node degree of 0.75, an average local clustering coefficient of 0.184, and an overall PPI enrichment p-value of 1.28e-11. These additional 236 inferred genes of interest are related to the differentially expressed genes identified in the IMFvsFF contrast samples. Here we identified 13 functional clusters (see corresponding subsection supplemental note for full cluster associations). The dominant clusters were enriched for DNA damage response and transcriptional regulation, including genes involved in DNA checkpoint control, non-homologous end joining, spliceosomal assembly, and mediator complex components. Additional clusters reflected lipid and energy metabolism, encompassing fatty acid degradation and synthesis, acyl-CoA and tricarboxylic acid (TCA) cycle enzymes, and 4-coumarate-CoA ligaseassociated acyl-chain remodeling. Several smaller clusters represented hormone-mediated-developmental signaling and post-translational processes, including MAPK and ethylene-activated signaling, protein tagging, glycosphingolipid biosynthesis, redox metabolism (NAD(P)H oxidase activity), gibberellic-acid signaling, and macromolecule depalmitoylation.

## Discussion

Although sex determination in cannabis is primarily genetic, phytohormone signaling exerts a strong modulatory influence on sex expression^7,25^. Adal et al. (2021) provided a broad overview of how hormones such as abscisic acid, auxin, cytokinin, ethylene, and gibberellin contribute to sex expression, consistent with our findings. Our results further suggest that these phytohormone transduction pathways are intricately connected to lipid and membrane metabolism, cell wall biogenesis, and embryogenic development, presumably underlying the wide range of phenotypic differences between male and female plants. Consistent with previous transcriptomic analyses^7,25,36,53^, genes differentially expressed in our IMF vs FF contrasts include those involved in seed, pollen, and flower development, ethylene signaling, and phytohormone response pathways (Figure 4). The parallels between our results and those of Adal et al. (2021) are substantial, despite differences in plant genotypes and masculinizing agents (Figure 2). This suggests a shared core set of gene regulatory networks governing transcriptional responses associated with sexual plasticity in cannabis. Notably, this consistency emerged even though our study utilized colloidal silver, while Adal et al. (2021) employed silver thiosulfate. The overlap and divergence in the total number of differentially expressed genes across both datasets (Table 1) further indicates that silver thiosulfate may be the more potent masculinizing agent; however, colloidal silver was selected in our study due to its safer handling and compatibility with available laboratory facilities.

**Figure 4.**
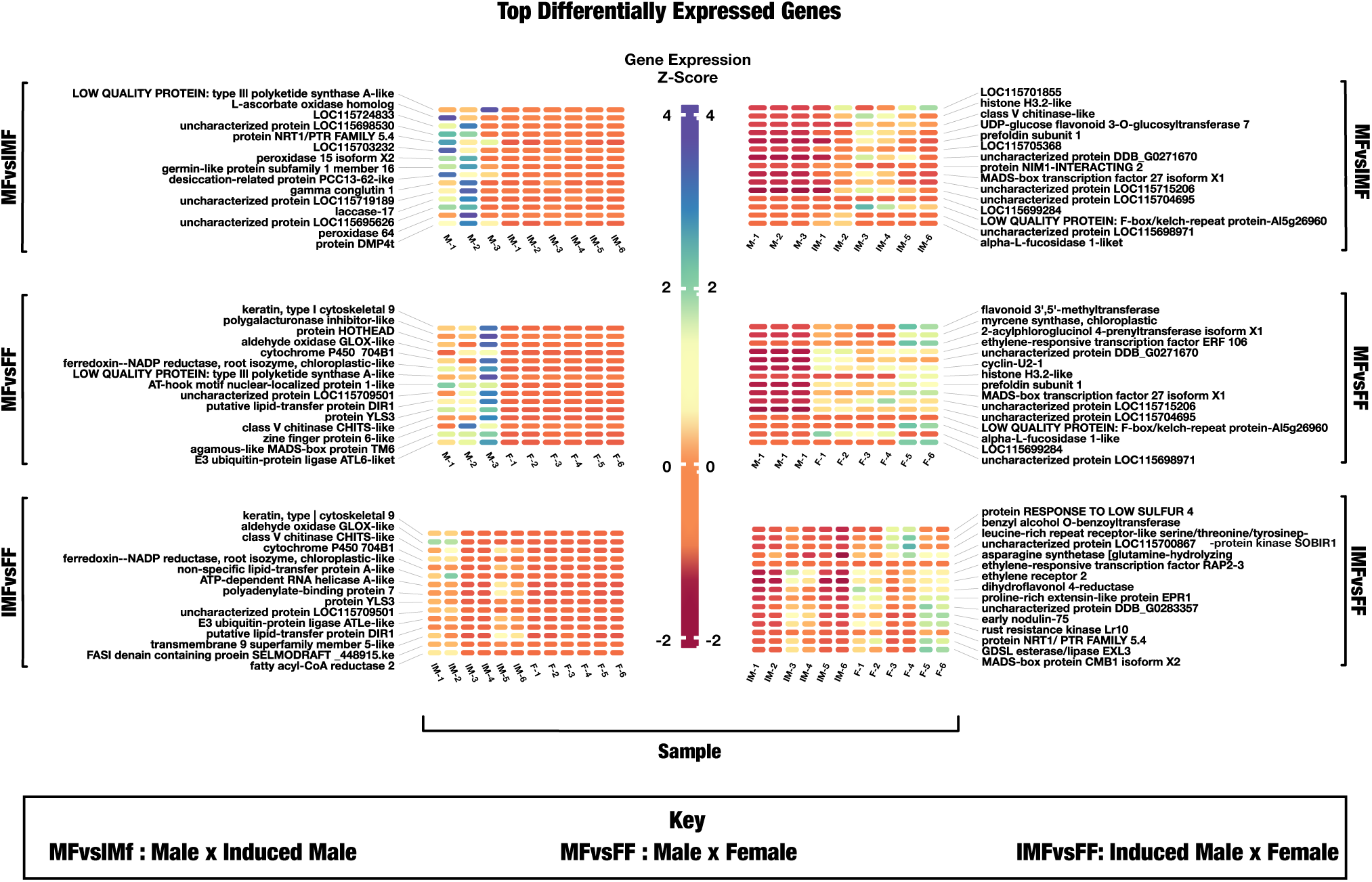
Heatmaps of differentially expressed genes across three contrasts. Z-score normalized gene expression values are shown for the top differentially expressed genes (DEGs) identified in pairwise comparisons: MFvsIMF: male flower vs. induced male flower, MFvsFF: male flower vs. female flower, and IMFvsFF: induced male flower vs. female flower. Heat maps on the left showcase genes up-regulated in the first treatment of each contrast (MF in the top row, MF in the middle, and IMF on the bottom) compared to the second treatment of the same contrast (IMF in the top row, FF in the middle, and FF on the bottom). The heat maps on the right show the up-regulation of the second treatment, compared to the first treatment, in each contrast. Protein annotations were derived using the Cs10G2P file and ontology inference. Substantial up-regulation of genes related to pollen development and phytohormone signaling in IMF samples suggests cross-talk between stress-response hormones. Expression levels in FF and IMF samples (IMFvsFF) indicate conservation of FF expression patterns in IMFs.

**Table 1.** Counts of unidirectional DEGs (log-fold change 1, p-adj 0.05) identified as overlapping across datasets post independent processing of both the Adal et al. 2021 and our RNA-seq samples into counts using Jack and identical differential expression analysis parameters via DESeq2.

|  | CN1 |  | CN2 |  | CN3 |  |
| --- | --- | --- | --- | --- | --- | --- |
|  | Up | Down | Up | Down | Up | Down |
| **Orozco** | **790** | **2,534** | **1,022** | **2,482** | **123** | **10** |
| **Orozco and Adal** | **69** | **56** | **1,002** | **6** | **1** | **0** |
| **Adal** | **1,753** | **494** | **19,973** | **41** | **744** | **1,689** |

**Table 2.** Total number of nodes and edges in the co-expression and PPI networks before and after filtration. Co-expression is entirely unfiltered, Co-expression Alpha 0.3 uses filtration factor alpha of 0.3 for preliminary filtration. Filtered Co-expression and Filtered PPI truncates nodes and corresponding edges which are not present in both the co-expression and PPI networks to create the Bipartite Network. Nodes and edges in bipartite network included for posterity. Compute significance scores (alpha) for weighted edges in a NetworkX graph as defined in Serrano et al. 2009, The resulting graph contains only edges that survived from the filtering with the alpha_t threshold. Nota performs a cut of the graph previously filtered through a proprietary function. PPI network was only filtered to match the cs10G2P 1-to-1 map.

|  | Number of Nodes | Number of Edges |
| --- | --- | --- |
| **Co-Expression** | 22,658 | 30,455,297 |
| **Co-Expression Alpha 0.3** | 22,630 | 3,431,798 |
| **Filtered Co-Expression** | 12,873 | 1,360,160 |
| **PPI** | 23,704 | 3,285,936 |
| **Filtered PPI** | 12,873 | 1,275,017 |
| **Bipartite Network** | 25,746 | 12,873 |

Similar to our dataset, Monthony et al. (2024) examined ethylene-related genes (CsERGs) associated with sex determination and sexual plasticity using transcriptomic data. Their classification of CsERG expression into Karyotype Concordant (KC), Floral Organ Concordant (FOC), and Unique Ethylene-Related Gene (uERG) patterns provides a refined view of ethylenes role in reproductive differentiation. They showed that FOC genes maintain consistent expression across individuals sharing floral organ type despite differing karyotypes, whereas uERGs vary across FF, MF, and IMF phenotypes, indicating a complex regulatory interplay.

Our findings complement and expand upon this framework by revealing broader phytohormone-mediated contributions to sex determination. Specifically, our results suggest that silver ion treatment promotes masculinization by inhibiting ethylene signaling through the disruption of metal ion homeostasis. This interference appears to trigger a cascade of transcriptional and metabolic reprogramming events that facilitate embryogenic differentiation and the re-establishment of male floral identity.

Further supporting this, García-de Heer et al. (2024) suggest that hormone pathway manipulation can induce sex reversion, with monoecy in cannabis likely being linked to spatial variability in sex-influencing hormones, which act upstream to repress or promote floral identity genes^15^. This aligns with our findings, reinforcing the idea that sex expression is tightly linked to flowering and influenced by both genetic and hormonal factors. Although C. sativa has a chromosomal system (XX for females and XY for males), environmental conditions and hormone fluctuations can override genetic determinants, leading to the formation of male flowers on genetically female plants.

Marker-assisted breeding, SNP mapping, and QTL analysis are increasingly employed to enhance seed production and fiber quality in commercial hemp cultivation^25^. Male-associated MADC2 PCR markers have been used to distinguish male phenotypes from dioecious and monoecious females, providing a genetic foundation for breeding programs^54,55^. However, these approaches rely on previously characterized molecular markers, which may not fully capture the regulatory complexity of sex determination in non-model species. In the following sections, we contextualize these differential expression and RWR analyses to explore the broader regulatory mechanisms underlying sexual plasticity in cannabis.

### Stress and Phytohormone Signaling Crosstalk

The induction of stress response genes was a common theme across all differential expression contrasts, with gene functions demonstrating that both natural sex differentiation and colloidal silver-induced sex reversal involve stress-related pathways as corroborated in relevant literature^32^. Aldehyde oxidase GLOX-like (LOC115709228), involved in detoxifying reactive aldehydes and its Arabidopsis t. homolog is known to be regulated by MYB80 transcription factor in the development of tapetum and pollen^56^, was up-regulated in both MFs (MFvsFF) and IMFs (MFvsIMF, IMFvsFF), indicating a potential role for oxidative stress response in general plant male development. Oxidative stress is a common consequence of environmental perturbations and developmental transitions, and its management is crucial for maintaining cellular homeostasis^57^. Up-regulation of genes like Callose synthase 5 (LOC115714911) in IMFs (IMFvsFF) and F-box/kelch-repeat protein (LOC115718742, At5g26960), in both IMFs (MFvsIMF) and females (MFvsFF) suggests that various stress response pathways may be activated concurrently during sex differentiation and in response to colloidal silver treatment (Figure 4). F-box/kelch-repeat protein (LOC115718742, At5g26960), which modulates proteasomal activity under stress conditions, is also involved in protein ubiquitination, a critical process for proteostasis under stress conditions^58^. The up-regulation of stress-related genes in IMFs likely reflects a stress-adaptive mechanism to regulate protein turnover during physiological adjustments induced by colloidal silver treatment or environmental signal cues.

In our analysis we observed the disruption of the sexual development of treated female cannabis, by a change in the metal ion homeostasis which directly affects the ethylene biosynthesis signaling pathway. However, it is known that the signaling pathways of phytohormone networks are highly interconnected, suggesting that gene members are shared between these networks (Figure 5)^27^. The role of ethylene in sex determination is well-documented in various plant species, and its disruption can lead to alterations in sex expression^59^. We identified several phytohormone related genes in our differential expression analysis in addition to genes in the ethylene signaling pathway. In our RWR inference set for MFvsIMF we identified several proteins related to the Giberellic, Jasmonic and Brassinosteroid mediated signaling pathways (XP_030496910 DELLA protein GAIP-B, XP_030501877 transcription factor MYC2, XP_030488074 somatic embryo-genesis receptor kinase 2). Consistent with our DEG results for MFvsIMF, several of the MFvsIMF RWR inferences clustered to hormonal and stress-response pathways (Induced systemic resistance, Jasmonic acid and salicylic acid signaling pathways, Regulation of Jasmonic acid signaling and ubiquitin-like conjugation). These results elucidate the possible molecular mechanisms that cannabis employs during transcriptional regulation and phytohormone signaling to adaptively mitigate cellular stress and regulate reproductive organ development under abiotic stress conditions. One possibility is that abiotic stress acts to modulate the genetic architecture of sexual plasticity, leading genetically XX Females to produce male flowers, perhaps as an evolutionary adaptation to ensure seed production^10,16,17^. However, further functional experimentation is required to validate these predicted interactions between major phytohormone gene networks, and their effects on sexual phenotypes.

**Figure 5.**
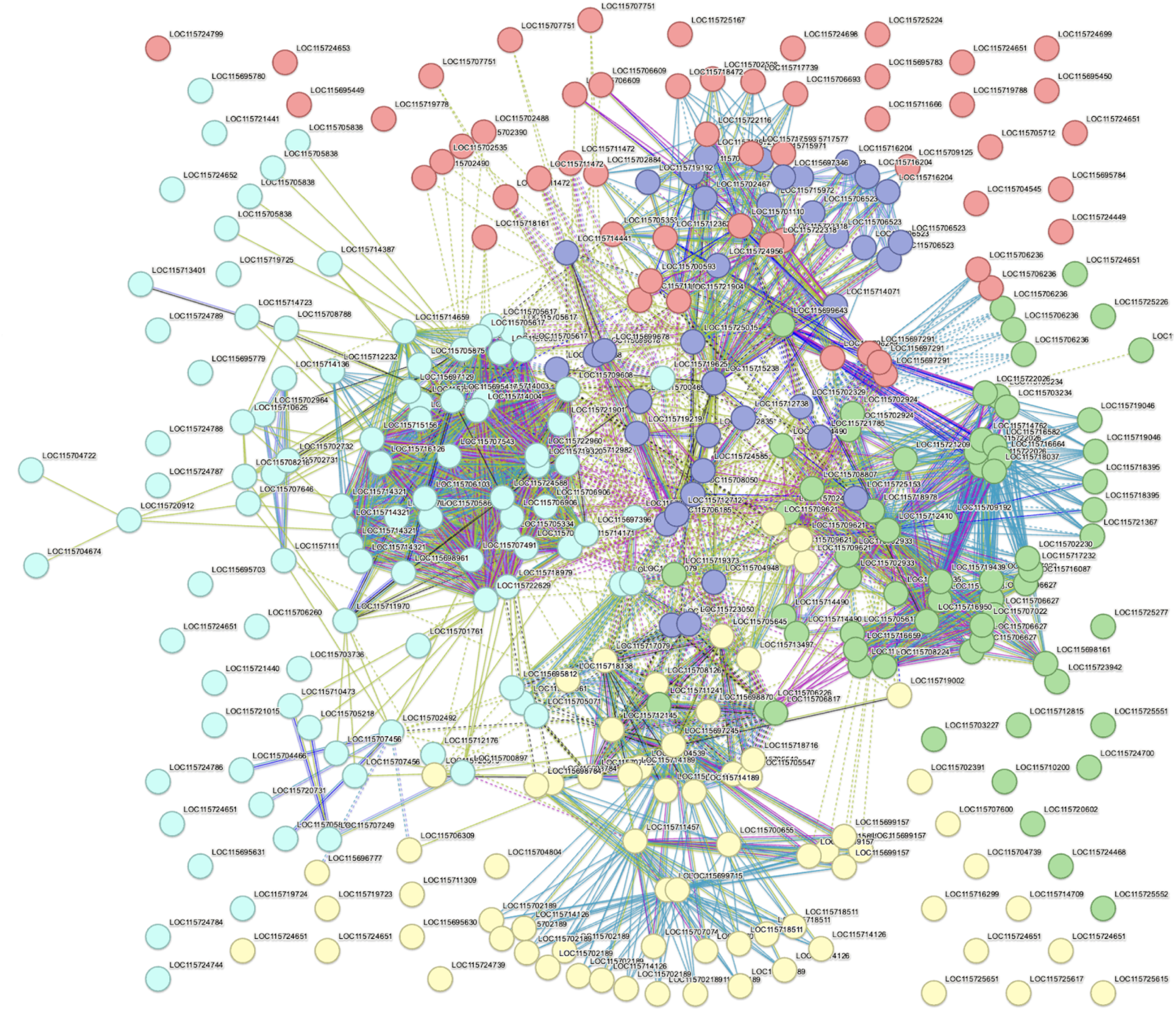
Protein-Protein Interaction Network between Five Major Phytohormone signaling pathways in cannabis (reference genome cs10). Green nodes: Abscisic acid signaling pathway, Yellow nodes: Jasmonic acid signaling pathway, Teal nodes: Auxin signaling pathway, Purple nodes: Ethylene/Brassinosteroid signaling pathway, Red nodes: Cytokinin signaling pathway. Edges are color-coded based on the type of interaction: Blue edges represent known interactions from curated databases, Magenta edges indicate experimentally determined interactions, Green edges signify predicted interactions from gene neighborhood, Red edges denote gene fusions, and Blue edges signify gene co-occurrence. Black edges represent co-expression, Yellow edges indicate text mining, and Light Blue edges denote protein homology. Dashed lines often represent inter-cluster connections, indicating relationships or interactions between different clusters (K-means 5 clustering applied).

More specifically, across all contrasts, genes directly and indirectly related to phytohormone signaling pathways: ethylene (ETH), abscisic (ABA), jasmonic (JA), auxin (AX), and gibberellin (GB) were differentially expressed. Cytochrome P450 704B1 (LOC115700892), involved in gibberellin biosynthesis, was consistently up-regulated in both true males (MFvsFF) and IMFs (IMFvsFF), supporting the role of gibberellins in promoting male reproductive development^60^. This aligns with studies in other plant species in which gibberellins have been shown to influence floral sex differentiation and promote the development of male reproductive structures^60^. MYB101 transcription factor (LOC115699028) was up-regulated in MFs but down-regulated in IMFs. MYB101, regulates pollen tube-synergid interactions during fertilization^61^ and has been linked in *Arabidopsis thaliana* to ABA-mediated stress responses, such as stomatal closure and transcriptional regulation^62,63^. Its differential expression further underscores the disruption of phytohormone signaling in IMFs caused by colloidal silver treatment. The categorization of clusters in the IMFvsFF RWR inferences continued to highlight the functional categories identified in the corresponding DEG results that highlighted key molecular processes differentiating IMF cannabis tissue from true female cannabis tissue. Our RWR inferences corroborate our DEG results and provide additional insight into the regulatory, metabolic, and stress-response pathways involved in the masculinization process.

Ethylene-responsive transcription factor ERF106 showed a complex pattern, being up-regulated in IMFs (MFvsIMF, IMFvsFF) but down-regulated in MF (MFvsFF) compared to FF. This suggests that colloidal silver treatment may disrupt ethylene signaling, potentially contributing to the induction of male traits^64^. The differential expression of genes functionally linked to metal ion transport and homeostasis, such as myrcene synthase (LOC115716063) and metal-nicotianamine transporter YSL7 (LOC115701423) in the MFvsFF contrast, further highlights the potential role of metal ions and metal ion homeostasis in secondary metabolism and sex differentiation in response to disruption of the interconnected phytohormone gene network crosstalk. Metal ions play essential roles in various biological processes, and their homeostasis is crucial for proper plant development and stress response^65^.

Taken together, the findings of each of our DEG contrasts thus demonstrate that up-regulation of stress-responsive, phytohormone-regulated, and developmental genes in IMFs reflects an adaptive transcriptional landscape shaped by colloidal silver treatment. The interplay between the ethylene, GA, and JA signaling networks appears to mediate the genetic reprogramming that underlies sexual plasticity.

### Metal Ion Homeostasis and Cross-Cellular Communication

The disruption of ethylene signaling by silver ions (Ag^+^), (Figure 6), not only inhibits ethylene receptor activity but also appears to disrupt the regulation of key proteins associated with oxidative stress and lignin metabolism. For example, expression of the blue copper protein (LOC115699453), was up-regulated in MFs but down-regulated in IMFs, likely due to incomplete enzymatic activity of the ethylene receptors, being blocked by the applied silver ions (Figure 6)^33^. IMFs exhibited down-regulated expression of genes linked to auxin signaling and desiccation responses, such as desiccation-related protein PCC13-62-like (LOC115725418)^66^, reflecting stress-adaptive transcriptional programs that respond to oxidative stress and hormonal imbalance. Furthermore, the down-regulation of ethylene receptors (LOC115721785) and ethylene-responsive transcription factors (LOC115723033) in IMFs (IMFvsFF) reinforces the disruption of ethylene signaling pathway by colloidal silver. IAA-amino acid hydrolase ILR1-like 4 (LOC115707590), involved in auxin metabolism, was down-regulated in IMFs and up-regulated in FFs (IMFvsFF). Auxin has been reported as a key regulator of plant development, playing a role in sex differentiation^67,68^ thus indicating a potential role for auxin in the prioritization of male embryogenic tissue reprogramming in treated samples.

**Figure 6.**
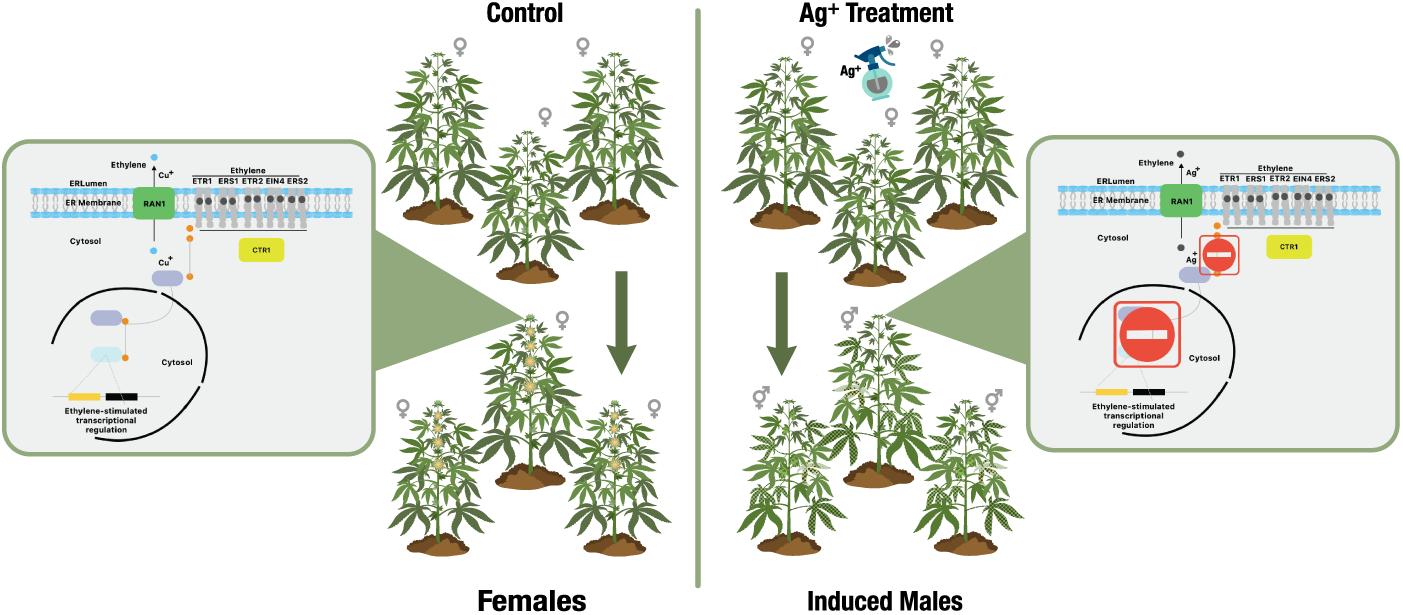
Colloidal Silver Interruption of Ethylene Signal Transduction: Silver ions (Ag^+^) inhibit ethylene (C_2_H_4_) signal transduction in cannabis by out-competing copper ions (Cu^2+^) at the ethylene receptor sites. This prevents receptor activation and reduces ethylene-related gene expression. The control panel shows uninterrupted ethylene signaling in female cannabis plants, while the treatment panel illustrates how Ag^+^-induced hormonal stress disrupts this process, enabling the monoecious female plant to leverage its sexual plasticity for seed production under stress.

From our differential expression analysis we observed interesting expression profiles of metal ion related genes such as the blue copper protein (LOC115699453) and probable metal-nicotianamine transporter YSL7 (LOC115701423) were up-regulated in our MFs but not in our IMF samples. These metal homeostasis related gene expression profiles suggest that our treatment disrupted metal ion homeostasis in our treated samples. Within our MFvsIMF RWR inference set we observed 40 genes associated with metal ion binding ontology. Cytochrome P450 CYP73A100 (XP_030493605) was an inferred gene related to the findings of our MFvsIMF DEG’s, the cytochrome P450 enzyme family are known to play crucial roles in the biosynthesis of secondary metabolism, defense and response to environmental stressors^69^. This observation is interesting as it seems that metal ion related genes from both the differential expression analysis and the RWR analysis for the comparison of MFs and FFs against our IMF samples have persisted. Our findings highlight the potential role of metal ions and metal ion homeostasis in secondary metabolism and sex differentiation in response to disruption of the interconnected phytohormone gene network crosstalk. The application of silver ions to plant tissue is known to disrupt the ethylene signaling pathway^35^ which is likely due to transcriptomic compensation to satiate incomplete enzymatic activity of the ethylene receptors dependent on copper ion binding, being blocked by the applied silver ions^70–81^.

### Multi-layered Regulation of Sex-Associated Genes

Transcription factors play a central role in orchestrating the complex gene expression changes associated with sex differentiation and the response to colloidal silver. Agamous-like MADS-box protein TM6 (LOC115714657) was up-regulated in MFs (MFvsFF), while MADS-box transcription factor 27 isoform X1 (LOC115704295) was up-regulated in IMFs (MFvsIMF, IMFvsFF) and FFs (MFvsFF) and down-regulated in MFs (MFvsIMF, MFvsFF). These findings complement the role that MADS-box transcription factors play in regulating distinct aspects of female floral development. MADS-box genes are well-known for their roles in floral organ identity and flowering time regulation, and their differential expression can considerably impact plant development^82^. Transcription factor bHLH91 (LOC115713370) was up-regulated in IMFs (IMFvsFF), as were transcription factor ABORTED MICROSPORES (LOC115706375), and PHD finger protein MALE MEIOCYTE DEATH 1 (LOC115721367), a probable transcription factor required for chromosome organization and progression during male meiosis^83^. Both of these transcription factors were down-regulated in FFs (IMFvsFF) and up-regulated in IMFs, highlighting their potential roles in male reproductive development and the response to colloidal silver in cannabis. Fittingly, the high transcriptional re-configuration observed in our samples was purportedly used to shift our samples cell wall and lipid metabolism. The MFvsFF contrast RWR inferences provide additional key differences between male and female cannabis at the metabolic, regulatory, and structural levels(See CN2_annotated_multiplex_2 in RWR Results and Multiplex Files supplemental data folder online).

Genes involved in protein stability (ubiquitination) folding and degradation were differentially expressed, indicating a potential role for proteostasis in sex differentiation and response to colloidal silver. E3 ubiquitin-protein ligase ATL6-like (LOC115718105) was up-regulated in MFs (MFvsFF) and IMFs (IMFvsFF), suggesting a role for protein degradation in male development. MFs exhibited lower expression of ubiquitination-related genes, including F-box protein SKIP14 (LOC115699558), which is also associated with negative regulation of ABA signaling^84^. The ubiquitin-proteasome system is a key regulator of protein turnover and plays a crucial role in various cellular processes; hormone signaling, the regulation of chromatin structure and transcription, tailoring morphogenesis, responses to environmental challenges, self recognition and battling pathogens^85^. The prefoldin chaperone complex, of which Prefoldin subunit 1 (LOC115705604) is a subunit, is a conserved hetero-hexameric co-chaperone that delivers nascent polypeptides to the TRiC/CCT chaperonin for folding, thereby maintaining proteome integrity and cellular homeostasis^86–88^. Prefoldin subunit 1 (LOC115705604) was up-regulated in IMFs (MFvsIMF) and down-regulated in MF (MFvsFF) suggesting a potential role for chaperone complex related proteins in the response to colloidal silver and sex differentiation in cannabis. The malfunction of the pre-folding complex has been associated with altered growth and environmental responses^89^. These findings suggest that proteostasis is involved in the molecular mechanism driving sexual development, an interesting pairing with the high transcription factor modularity that has been sexual development.

### Cell Wall and Lipid Remodeling in Reproductive Development

Changes in cell wall-related genes were observed across the differential expression contrasts. Polygalacturonase inhibitor-like (LOC115700758) was up-regulated in both MFs (MFvsFF) and IMFs (IMFvsFF), suggesting a role for pectin modification in male reproductive development. Cell wall modifications are essential for plant growth and development, and alterations in cell wall composition can influence various physiological processes including plant morphogenesis^90,91^. The protein HOTHEAD (LOC115717125), involved in cuticle development, was up-regulated in MFs (MFvsFF), while alpha-L-fucosidase 1-like (LOC115709178), a lysosomal enzyme that breaks down fucose-containing glycoproteins and glycolipids^92,93^, was down-regulated in MFs (MFvsFF) and up-regulated in FFs. These additional findings suggest that different cell wall modifications may be associated with male and female cannabis morphogenesis. Similarly, Laccase-17 (LOC115719496) and Peroxidase 64 (LOC115723402), are both implicated in lignin metabolism, and showed up-regulated expression in MFs while being down-regulated in IMFs. Laccase-17 and Peroxidase 64 are involved in lignin degradation, detoxification, and oxidative stress responses^94,95^, suggesting that IMFs may be actively modifying lignin development as part of the physiological shift post-treatment. This reduction in lignin-related gene expression in IMFs highlights potential metabolic cost associated with transitioning from female to male reproductive structures.

We observed up-regulation of cell wall and lipid metabolism/transport-related genes in MF samples compared to FF and IMF samples (Figure 4). In our differential expression results for MFvsFF we found two lipid transport proteins (LOC115718067 putative lipid-transfer protein DIR1, LOC115707560 non-specific lipid-transfer protein A-like) were up-regulated in MF tissue samples while being down-regulated in FF tissue samples. Additionally in our MFvsIMF RWR inferences we identified a gene involved in phytohormone signaling but also clustered to di-terpenoid biosynthesis and cellular response to lipid metabolism (XP_030496910 DELLA protein GAIP-B) and protein XP_030487533 ABC transporter G family member 9 which is linked to ABC transporters in lipid homeostasis (MAP-1369062). Also protein (XP_030497001 REF/SRPP-like protein At1g67360) that is directly involved in lipid droplet formation, and clustered to proteins associated with the regulation of fertilization, seed dormancy process and pollen germination. These results support the idea that lipid-based signaling may be sex-specific in reproductive tissue differentiation^96–102^.

Several of our MFvsIMF RWR network inferences were ontologically associated to both primary cell wall biogenesis and embryo-genesis related terms. COBRA-like protein 6 (XP_030478181) was another inferred MFvsIMF related protein clustered to proteins involved in Plant-type primary cell wall biogenesis. Universal stress protein PHOS34 (XP_030483164) and kiwellin-like (XP_030480936) clustered to anther development and spermatogenesis, respectively. Additionally, somatic embryo-genesis receptor kinase 2 protein (XP_030488074) clustered to receptor serine/threonine kinase binding and flower morphogenesis. We also observed proteins associated with cellular component assembly involved in morphogenesis and pollen wall exine assembly in our MFvsFF set. Our IMFvsFF inference set also had several proteins associated to spliceosomal snRNP assembly, Glycosphingolipid biosynthesis, Ubiquitin mediated proteolysis, MAPK signaling pathway in plants, further emphasizing the interplay between lipid metabolism, signal transduction and reproductive development. These findings are consistent with previous studies linking COBRA-like proteins to cellulose deposition and primary cell biogenesis, processes fundamental to morphogenesis and growth^103,104^. Likewise the identification of somatic embryo-genesis receptor kinase 2 protein (XP_030488074) of somatic embryogenesis receptor-mediated signaling that governs floral and tapetal development^105–107^. The presence of proteins associated with pollen wall formation and cellular morphogenesis further reflects conserved mechanisms in male gametogenesis^108,109^. Additionally, the enrichment of splicesomal snRNP assembly, glycosphingolipid biosynthesis, and MAPK signaling components among IMFvsFF inferences highlights how post-transcriptional regulation, lipid metabolism, and signal transduction collectively modulate reproductive development^110–112^.

Intriguingly, the lipid-mediated cellular activity identified here draws clear parallels with mechanisms described in mammalian systems where Hedgehog signaling pathway ligands ( Sonic Hedgehog, Indian Hedgehog, and Desert Hedgehog) exert developmental signaling by associating with membrane micro-domains (lipid rafts) and undergoing proteolytic and lipid modifications during secretion. In mammals, secreted morphogens can bind to lipid rafts on the surface of secreting cells. These rafts, with their characteristic glycosphingolipids and serine/tyrosine protein receptor composition, facilitate signaling that influences development in nearby cells. Alternatively, they can be released via proteolysis or packaged into vesicles or lipoprotein particles to act on distant cells^113,114^. In plants, lipid rafts have also been proposed as functional membrane micro-domains that compartmentalize signaling and trafficking, as reviewed in Solonaceae species including tomato^115,116^. Moreover, recent work in tomato has demonstrated that membrane-bound sterols and glycosylated sterol derivatives, components of putative raft domains are dynamically regulated during development and stress responses^117^.

The parallels between these mammalian and plant lipid-micro-domain systems suggest that in *Cannabis s.*, lipid rafts may similarly act as hubs for intercellular or long-distance signaling, particularly during sex differentiation. Our data support the hypothesis that lipid-rich micro-domains could mediate cross-cellular communication in embryo-genic or flowering tissue, facilitating the reprogramming of such tissue in response to environmental stress signals. This hypothesis is further bolstered by observations in tomato glandular trichomes, whereby epidermal structures produce specialized metabolites in coordination with lipid-based processes, phytohormone signaling, and stress-responsive transcriptional networks^18,24,118^. In both systems trichomes in cannabis and epidermal/secretory cells in tomato the overlap of phytohormone, lipid-metabolism and stress networks reinforces the potential role for lipid-domain-mediated signaling in reproductive tissue plasticity.

A key observation was the reprogramming of embryogenic tissue development through long-range intracellular cascade signaling, primarily mediated by lipid metabolism. The introduction of high-affinity silver ions through colloidal silver treatment disrupted metal ion homeostasis and phytohormone network cross-signaling, triggering widespread transcriptional and metabolic reprogramming. Notably, treated flowers exhibited gene expression changes related to lipid-protein transport, lipid metabolism, and cell wall development/modification. This physiological shift, coupled with hormonal imbalance, initiated a cascade of cross-cellular signaling, ultimately leading to extensive protein ubiquitination and proteolysis. Our analyses identified genes associated with cell wall morphogenesis and embryogenic tissue homeostasis, indicating that silver ion treatment perturbs phytohormone signal transduction through metal ion imbalance. This disruption, mediated through lipid-domain signaling, appears to drive a coordinated reprogramming of hormonal, metabolic, and structural pathways underlying sexual plasticity in*Cannabis s.*.

Capturing this level of cross-cellular complexity required analytical approaches capable of integrating transcriptomic and proteomic interactions across multiple biological layers. The network inference framework implemented through Nota, in tandem with the data accessibility capabilities of Jack, provides the computational foundation for this systems-level interpretation and represents a significant step forward in the functional annotation of complex traits in non-model species.

### Advancing Complex Trait Analysis and Enrichment with Jack and Nota

A key innovation in this study was the application of Nota for gene network annotation in cannabis. Unlike previous studies that align cannabis transcriptomic data to closely related model organisms, we used a multilayer biological network inference approach. Our method integrated high-order interactions and functional annotations derived from a species-representative proteome via the STRING database, providing a more tailored and robust framework for gene annotation in non-model organisms. We suggest that readers search for all functional terms of interest with our Cannabis Species Representative STRING PPI.

Our findings suggest that Nota offers a flexible alternative for functional annotation, particularly for systems with limited reference genomes or established molecular markers. By incorporating high-order interactions, Nota improves on traditional annotation methods by capturing complex regulatory networks rather than relying solely on primary sequence homology. We encourage the transcriptomics community to further explore functional annotation through network-based approaches, which can enhance the resolution of biological inference in non-model species like cannabis.

For further exploration of phytohormone network clustering, visit the STRINGDB synthetic proteomes linked in each respective contrast section within the results. These network analyses incorporate various data types, including gene expression and PPI networks, to make predictions about individual gene function and the relatedness of complex trait architectures. This innovative platform allowed us to connect transcriptomic evidence with inferred relationships in the phytohormone signaling network, as demonstrated by the highly interconnected protein-protein interaction network of the reference cannabis genome (cs10) (Figure 5). This is achieved through Nota’s use of multilayer network-based methods, enabling the identification of gene modules and community structures that relate to complex traits more efficiently.

Nota’s ability to identify higher-order interactions between transcriptomic and proteomic datasets was pivotal in our study. By integrating these datasets within a multilayer framework, we were able to uncover the complex cross-talk mechanisms between different biological layers, providing a comprehensive view of the regulatory networks driving sexual plasticity in cannabis. This approach uncovered novel relationships, such as the clustering of stress-responsive and developmental genes, which may otherwise have been overlooked. Nota can incorporate various (gene co-expression and PPI) layers into a bipartite coupled multiplex network to infer additional functional annotation of complex trait architectures. For example, we visualized the gene co-expression layer using a filtered network representing a 100,000-edge sample of the 1.36 million-edge network (Figure 7), laid out with the integrated ForceAtlas2 spring layout algorithm to reveal key community structures.

**Figure 7.**
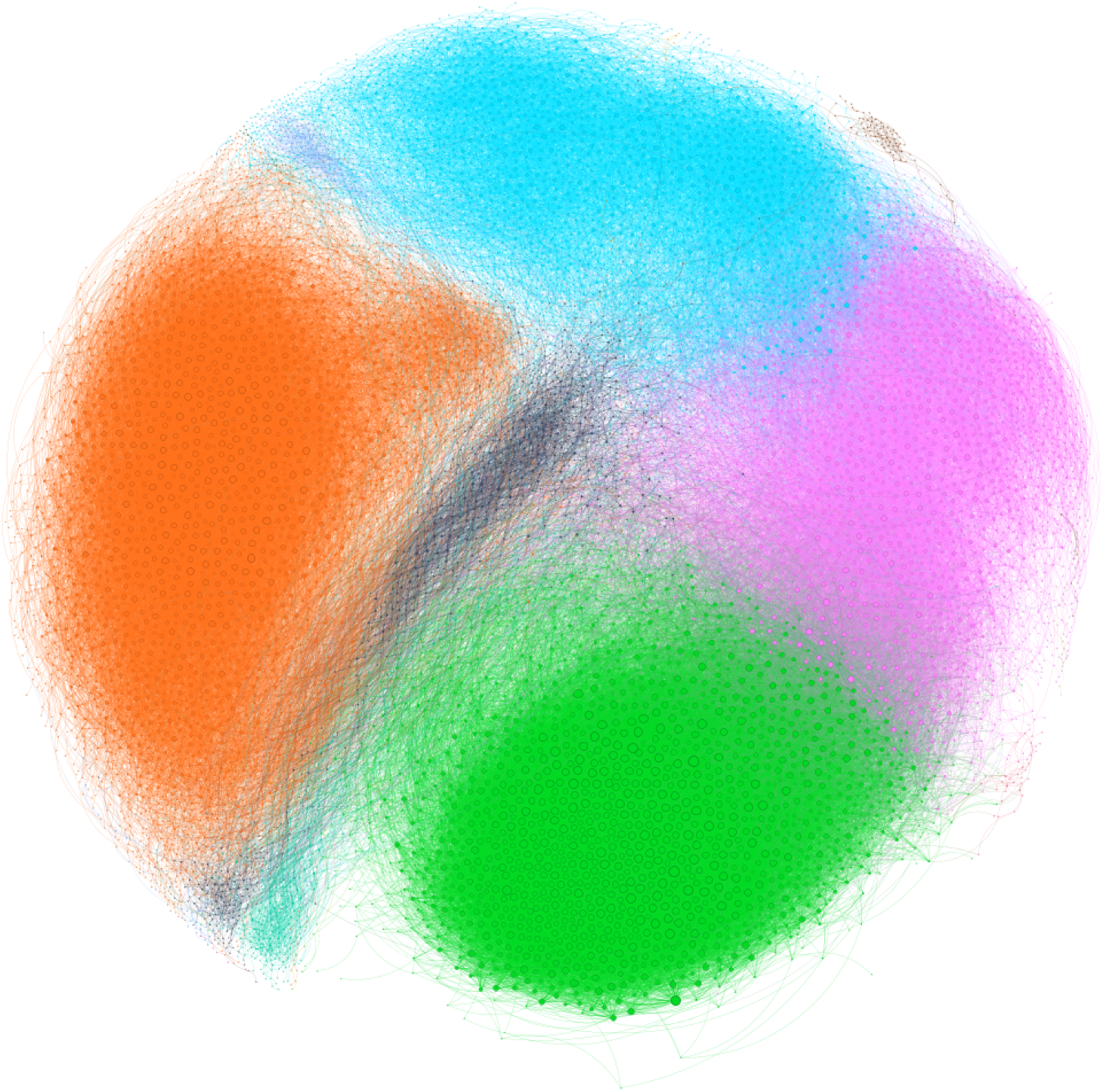
Filtered Co-expression Gene Network. A 100,000 edge sample of the 1.36 million edge gene co-expression network (11,533 nodes) laid out using the ForceAtlas2^129^ spring layout algorithm. Nodes and edges are colored by community membership. The top five communities account for 96.92% of identified comminutes detected within our confidence threshold. The pink community represents 26.42% of nodes were enriched to photosynthesis. The green community represents 23.79% of nodes that were enriched for translation. The blue community represents 19.72% of nodes that were enriched to regulation of active regulation of intracellular localization and trafficking of macro molecules GO terms. The black community represents 13.62% of nodes that were enriched for vesicle mediated transport GO terms. The orange community represents 8.12% of nodes that were to plant type cell wall organization and biogenesis GO terms.

Nota enables users to efficiently analyze gene expression data through its integration of advanced bioinformatics pipelines, and Jack functions as a user-friendly Google Gemini-powered bioinformatic analysis assistant. Traditional gene discovery workflows often require months of data analysis, model building, and validation. By leveraging network inference and bioinformatic tools, Nota considerably reduces the time researchers spend searching for genes associated with complex traits, without sacrificing the rigor of traditional genetic analysis. Nota empowers users to prioritize experimentally validated genes of interest, facilitating a deeper and more precise annotation of gene network architectures for complex traits requiring only novice coding literacy. This accessibility is transformative for geneticists, molecular biologists, and plant and animal breeders who employ functional annotation tools which rely solely on pair-wise interactions for analyzing and annotating gene expression data and identifying candidate genes.

The goal of Jack is to combine the distributed data infrastructures within the Bioinformatics community to both increase researcher efficiency and data accessibility to the public, we will be releasing a public version for these capabilities in the future. The infrastructure for data aggregation is present through multiple government institutions, however, is not presented in a way which is easy to access for multiple stakeholders. Further development is guided by principles to increase accessibility to public resources through data stewardship^119^. In addition, the functionalities of Jack support the long-term data preservation between multiple entities, to support researchers in better data practices overall and continue to support open-science data processes. Increasing accessibility in such a way is beneficial for multiple facets of the socio-economic sphere as found by^119,120^. Jack decreases the technical barriers to this open data access, thus decreasing the overall financial cost of researchers who do not reside in the global north.

### Conclusions

Across all differential expression and RWR results, the transcriptional landscape revealed notable differences in genes associated with phytohormones, metal ion homeostasis, lipid metabolism, cell wall remodeling, transcriptional regulation, stress responses, and both primary and secondary metabolism. The MFvsIMF contrast highlighted the disrupted reproductive function and stress adaptation in IMFs, while MFvsFF underscored the distinct hormonal and structural pathways differentiating male and female reproductive strategies. The IMFvsFF contrast revealed the transcriptional cost and incomplete physiological transition from FF to IMF phenotypes. These findings support our hypothesis that several systems respond to environmental and hormonal signals to induce sexual plasticity in cannabis inflorescence tissue.

These functional trends in our interpretation of our results are reflected in the structural layout of the co-expression network (Figure 7). The network’s modular organization follows the biological hierarchy described in the figure caption, with each community representing a functionally cohesive set of coexpressed genes whose shared regulation suggests participation in distinct yet interconnected cellular programs. Together, these modules capture the coordinated transcriptional architecture underlying the physiological response of colloidal-silver treated female cannabis flower tissue. The network topology supports a model in which environmental perturbation drives the cross-cellular communication of metabolic and regulatory pathways via phytohormones signal transduction, metal homeostasis and cellular remodeling in masculinized cannabis.

The DEG and RWR analyses reveal potentially novel functional relationships that warrant further experimental validation. Although network construction parameters and interaction thresholds were rigorously defined to ensure robust inference, confirming the predicted gene and protein functions through targeted assays remains an important next step. Our representative proteome is a demonstration of new methods for improved annotation and gene ontology within cannabis without needing to develop new model organisms. Despite challenges such as heterogeneous environments, varying genome assembly quality, and legal constraints, integrating genomic, transcriptomic, and inferred functional proteomic data, combined with network inference, advances our understanding of *Cannabis sativa* sex determination. By unraveling the cross-cellular communication underlying this process, we establish a framework for targeted modulation of sex expression via marker-assisted breeding systems or chemical manipulation of gene expression.

## Methods

### Plant Growth and Treatment

All cannabis material for this analysis was identified and processed by and on the premises of Skinny Pineapple Inc, Boulder Colorado, in accordance with local and state laws and regulations. No specimens from this study were preserved as specimen vouchers in herbarium. The three crosses selected for this study contain the same paternal parent Sour Strawberry, which had been crossed with Super Lemon Haze (SxSS), Flaming Cookies (FCxSS), and Golden Goat (GGxSS). These seeds were grown to produce four clones, one of which was pre-maturely flowered to ensure the sex of the plant was female, while the other three clones were grown to maturity under isolated and standardized growing conditions. The controlled conditions included consistent watering, temperature, humidity, and nutrient schedules, as well as placement under the lights in the grow tent. The plants were systematically rotated around the tent every week to receive equal light and physical data were recorded each week. The plants were given 5.5 grams of VegBloom (Hydroponic Research, San Diego, California) per gallon of water, with an average of 10 gallons of water used to water the 15 plants (6 un-treated female, 3 male, and 6 treated female). To induce male flowers on these female plants, treatment branches on each plant were labeled and sprayed daily with 50 PPM colloidal silver (Silver Mountain Minerals, West Jordan, Utah) beginning the first day of flower and continuing until mature male flowers were produced on every plant. Treated plants were isolated to a separate growing environment to ensure un-treated female and male plants were not contaminated with colloidal silver. Subjectively, there appeared to be a high similarity in the early growth of the plants; however, as the plants grew, more noticeable differences developed.

### RNA Extraction, Library Preparation and Sequencing

RNA was extracted from 15 samples total: 6 female flowers, 6 IM flowers, and 3 male flowers. The plant material was cut from the plant and immediately stored in a ribonuclease (RNase) inhibitory solution to prevent RNA degradation. Extraction was then completed using the RNeasy mini kit (Qiagen, Redwood City, CA) following the manufacturers protocol for using the RLC buffer. This protocol extracted total RNA libraries from the treatment and control branches of each plant, including ribosomal RNA (rRNA), transfer RNA (tRNA), messenger RNA (mRNA), and other small RNAs. These RNA samples were stored at -80̌rC in the extraction buffer until library preparation began to prevent degradation.

These samples were sent to the University of Colorados BioFrontier Sequencing Core for preparation of libraries. Fifteen libraries were prepared between the fifteen selected individuals with two technical replicates of each control and treatment branch. BioFrontiers performed quality assessment using the BioAnalyzer/TapeStation and Qubit, mRNA isolation using Poly-A beads, and reverse-transcriptase (RT)-PCR to index the libraries for multiplex sequencing. These steps were performed using the NEXTflex Rapid Directional mRNA-Seq Kit Bundles (Bioo Scientific, Austin, Texas) according to the standard protocol included in the kit. To sequence these libraries, Illumnias NextSeq Sequencer (2X150, 300-cycle, paired-end reads) produced approximately 264 million reads between the 15 samples, with 17.6 million reads on average per library. See supplementary data file, CannaSeqReadMapSummary, online for comprehensive statistics on the libraries.

### Differential Expression Analysis

Raw paired-end RNA-seq libraries were generated for each of the 15 samples. Count files for each sample were then generated using Jack, Nota’s network inference assistive pipeline. Paired FASTQ files were interleaved into individual FASTQ files per sample and aligned to the cs10 reference genome index generated using the STAR: ultrafast universal RNA-seq aligner software^121^. The average percentage of uniquely mapped reads of our samples was calculated to be 73% with a mapping range of 60.8 - 78.1%. After generating BAM files for each sample via STAR, the BAM files were processed into a gene expression matrix using featureCounts: an efficient general-purpose program for assigning sequence reads to genomic features^122^.

Due to the complexity and user-specificity required for differential gene expression analysis, as well as a plethora of tools and resources available^123–125^, we maintained the statistical component of transcriptomic analysis independent from Nota. Our gene count sample profiles were analyzed using the DESeq2^123^ R package. The DESeq2 dataset was constructed by importing and integrating raw count data and sample metadata into a gene count matrix. Prior to differential expression analysis, genes exhibiting low expression (fewer than one count across all samples) were filtered out to remove non-expressed genes from our statistical inference. Differential expression analysis was conducted using DESeq2 functionality, which fits a negative binomial model to the gene count matrix. We specified the experimental design with the sex, ’ConditionType’, as a variable of interest to account for differences in condition types. Using our gene count matrix and sample metadata, we created three contrasts: male flower vs induced male flower, male flower vs female flower, and induced male flower vs female flower. Identical analysis was performed on the Orozco and Adal sample sets separate from one another because of the inconsistency of plant growth, treatment, and sampling methods used across the two experiments.

Statistical significance of differentially expressed genes was determined using an adjusted p-value (alpha) of less than 0.05 (p-adj 0.05) with an absolute log2 fold change thresholds of greater than 1 (log_2_FC |1|). These thresholds were selected for consistency with the differential expression profiling done in Adal et al. 2021. This filtering was done to focus on all expressed genes without bias towards up or down-regulated genes. Results were extracted and tabulated including log_2_FC values, p-values, and p-adj values. Normalized gene expression data were computed using DESeq2’s ’normalization’ function for subsequent analyses and visualization purposes. Expression data between the Orozco and Adal samples were filtered again using the same thresholds for p-adj values and log_2_FC values, then merged based on Locus IDs present in both sample sets for each respective contrast. Directionality of regulation was then determined based on the sign of the log_2_FC value for each gene and appended to the data frame. Using this directionality value for each gene, the datasets were compared and only genes with uniform directionality across datasets were conserved (Table 1).

Heat maps were generated to visualize the z-score transformed expression data of each sample to illustrate patterns of expression across different conditions (Figure4). Prior to this, these data containing Gene IDs, protein names, and regulation directionality values, for each contrast were then read into R and formatted for calculation of Z score. The expression value for each protein name was appended to the data frame and normalized using the R package dplyr^126^. Consistency of order, Z score range, and contrast specific samples were ensured within the script using dedicated functions. Heat maps were created using the package ComplexHeatmap^127^ with visual specifics, formatting, and color pallets produced by circlize^128^. The protein names displayed in heat maps were collected from the cs10 annotation file by parsing the protein name associated with each Locus ID.

### Network Construction

We employed Nota to curate and analyze a multilayer biological network (Figure 1 panel d) composed of a gene co-expression layer, a species-representative protein-protein interaction (PPI)^51^ layer, and a bipartite gene-to-protein annotation layer connecting them 1:1. The multilayer network we curated consists of two layers (Layer A and Layer B) of equal node cardinality but different edge structures. Layer A represents a filtered gene co-expression network constructed from RNA-seq reads as specified below. Subsequently, Layer B is the curated species representative protein-protein interaction network generated using STRINGDB as described below. Layers A and B are interconnected through a bipartite layer, where the bipartite nodes correspond to the union of nodes in Layer A and Layer B. In our network, the bipartite layer is representative of the previously identified correlation of encoding genes to translated proteins. The edges in the bipartite layer link nodes in Layer A to their associated counterparts in Layer B, facilitating inter-layer connectivity acting as the basis for further prediction.

### Gene Co-Expression Network

Due to our small sample size, we incorporated 15 samples from the Adal et al. 2021 cannabis masculinization transcriptomic analysis^25^. Post-hoc of our differential expression analysis, we pooled and processed all RNA-seq reads from our dataset and Adal’s. Using Jack we generated a multi-experiment gene count matrix from all 30 combined RNA-seq samples. Prior to the creation of the co-expression network, we removed all zero sum rows and normalized the gene count matrix. The co-expression network was then constructed using the Jack function which calculates the Pearson correlation coefficient between all pairs of genes in a gene count matrix. Given the size of the correlation matrix (generated from 30 samples and thousands of genes), Jack utilized multi-threading with the ThreadPoolExecutor module in Python. This enabled the processing of different gene-pair correlation calculations in parallel, considerably reducing the overall computation time by distributing the workload across multiple threads. Each thread processed a subset of gene pairs and identified significant correlations independently, which were then aggregated into the final co-expression network.

Jack constructed as an undirected graph object by using the python NetworkX library^130^, where nodes represent individual genes and edges represent significant co-expression relationships between gene pairs. Isolated nodes, which did not show significant co-expression with any other gene, were removed to improve the clarity and focus of the network. Once the correlation matrix was calculated, Jack applied a stringent filtering criterion by retaining only gene pairs with an absolute correlation value greater than or equal to 0.5, indicating a moderate to strong co-expression relationship. Gene pairs below this threshold were discarded to focus on biologically meaningful interactions. To fit the parameters which allows Nota to employ network inference across non-homogeneous mono and multiplex networks, we built a function within Jack that allows users to convert correlation matrices into distance matrices with a positive weight range. Additionally, we used Nota’s graph filtration function, which filters a weighted similarity matrix to generate an adjacency matrix, allowing for the extraction of significant connections in a graph. It applies a topological filtering method based on a paper by De Vico Fallani et al. 2017^131,132^. Summary network statistics, (nodes and edges pre and post filtering), were computed (Table2). The final network was saved in Graph Modeling Language (GML) format for subsequent analysis and visualization (Figure 7). We employed additional modularity-based community detection algorithms to visualize this large network that assign network members to communities based on link structure^46,133^.

### Species Representative Protein-Protein Interaction Network

Using Nota and Jack, we curated and filtered a PPI network for Cannabis generated using STRINGDB, a widely-used protein annotation tool and interaction network generator^51^. To ensure the accuracy of the network, we utilized a comprehensive but non-redundant proteome. We used Jack to construct this synthetic representative proteome from publicly available data and tools, using the Y chromosome from the BCMb Cannabis line (Salk Institute) and the Pink Pepper genome (RefSeq Accession GCF029168945.1) as reference material^134^. All available Cannabis protein sequences were downloaded from NCBI using Jack’s Gemini API functionality, excluding those submitted solely on the basis of in silico predictions from genome sequences.

To further refine the data, Jack reduced sequence redundancy by employing a two-stage alignment protocol using `miniprot`^134^. In the first stage, protein sequences were aligned to the reference genome, and overlapping alignments were parsed to retain a single representative protein sequence for each locus based on the highest chain score. In the second stage, Jack assigned representative multi-copy protein sequences to a single genomic locus using the highest chain score. These steps yielded a 1-to-1 representative proteome for Cannabis, which was used to generate the initial PPI network Cannabis Species Representative STRING PPI. Here again, we applied the correlation to distance matrix conversion function in Jack to maintain all the interaction scores.

To curate and filter the PPI network, we utilized the attribute of Nota that processes the PPI data generated by STRINGDB which incorporates preliminary filtering. Nota then converted the PPI network, using tools found in NetworkX, where nodes represent proteins and edges represent protein interactions, into a GML graph object. After filtering, isolated nodes (proteins without significant interactions) and reversed self-loop interactions (B-to-A) were removed by Nota to decrease the density of the network. The final network was exported in GML format for visualization, as well as a TSV edge list for further analysis. Summary statistics, including the number of nodes and edges in the filtered network, were saved to a TSV file as well (Table2).

### Bipartite Coupled Multilayer Network

By utilizing Nota to symmetrically connect the PPI layer and the Co-expression layer into a multilayer network, each node represents a gene or the derived proteins (using a Locus ID to protein ID key for cannabis). The gene and protein layers of the multilayer network are connected by a bipartite layer where each protein is connected to its encoding gene, this is done using the cs10 refrence genomes annotation file taking a 1:1 mapping for gene IDs to protein IDs. We generated a visualization of the bipartite network where Nodes are positioned using the ForceAtlas2 layout in Gephi, and plotted as layers using Matplotlib and NetworkX with code adapted from Klein^130,135,136^(see supplementary fig S1 online). Spring force layout is determined using the topology of the gene co-expression network, with the same positions repeated for the protein-protein interaction network.

### Network Ontology Transcript Annotation: Random-Walk-With-Restart

Random-walk-with-restart (RWR) is a powerful approach for exploring the topology of networks. By simulating the movement of a walker randomly traversing nodes through edges in a network, random walks are capable of capturing several structural properties of networks. In Nota’s RWR, the random walker, at each step, can navigate from one node to one of its neighbors or restart its walk from a node randomly sampled from a set of seed nodes. Nota, by enabling restart from one or several seed nodes, simulates a diffusion process in which the objective is to determine the steady state of an initial probability distribution. This steady state represents a measure of proximity between the seed(s) and all the network nodes. Here we used the top 60 differentially expressed genes in each experimental contrast and filtered these 180 genes to keep those which existed in the symmetric bipartite-coupled multilayer network as contrast-specific seed nodes (MFvsIMF: 30, MFvsFF: 36, FFvsIMF:37). Using RWR, Nota was able to predict genes which were related to the seed nodes through high order interactions. The inferred genes were then scored and mapped for further functional analysis.

### Jack

Annotated gene expression data were computed using the tabulated results from differential expression analysis for each contrast, filtered to retain significant data with p-adj 0.05 and log_2_FC |1|. Genes with the top 30 and bottom 30 log_2_FC values were designated as up-regulated and down-regulated, respectively. After this filtration, each Locus ID was cross-referenced with known cannabis gene and protein names for primary basic annotation, using annotation provided in the cs10 reference genome and using Jack’s Gemini and NCBI pubmed API functionality, we were able to generate a reliable list of available scientific journal references along with additional primary basic annotation for our cannabis transcripts and their respective encoded proteins. When a protein name was unavailable, we applied RWR, which predicted functional annotation by assigning a most-similar protein to an uncharacterized protein(s), and through sequential clustering to the Cannabis Species Representative STRING PPI now publicly available on STRINGDB. We were able to confidently assign function and, when available, ontology enrichment associated to its respective cluster.

## Data Availability

The RNA-seq data generated during the current study have been deposited on NCBI to the SRA under the bioproject PRJNA121253. The Adal et al. 2021 data analyzed in the current study can be found at NCBI SRA under the bioproject PRJNA66938. The datasets generated during and/or analysed during the current study are available in our *Figshare* repository, https://doi.org/10.6084/m9.figshare.30576740.

## Plant Material Processing Compliance

The authors processed the plant DNA, RNA, and other data in accordance with local and federal law, under the Research and Development permit 08 - 104227 to the Regents of the University of Colorado. All plant handling and identification was undertaken by C.P and K.W. on the presmises of Skinny Pineapple Inc. Boulder Colorado, following local and state law and regulation regarding the processing and handling of cannabis material.

